# Gephebase, a Database of Genotype-Phenotype Relationships for natural and domesticated variation in Eukaryotes

**DOI:** 10.1101/618371

**Authors:** Virginie Courtier-Orgogozo, Laurent Arnoult, Stéphane R. Prigent, Séverine Wiltgen, Arnaud Martin

**Affiliations:** Institut Jacques Monod, CNRS, UMR 7592, Université Paris Diderot, Paris, France; Institut de Systématique, Evolution, Biodiversité (ISYEB), UMR7205, CNRS-MNHN-UPMC-EPHE, PSL University, 45 rue Buffon, 75005 Paris, France; Department of Biological Sciences, The George Washington University, Washington, DC, USA

**Keywords:** genotype, phenotype, natural variation, domestication, animals, plants, comparative genetics

## Abstract

Gephebase is a manually-curated database compiling our accumulated knowledge of the genes and mutations that underlie natural, domesticated and experimental phenotypic variation in all Eukaryotes — mostly animals, plants and yeasts. Gephebase aims to compile studies where the genotype-phenotype association (based on linkage mapping, association mapping or a candidate gene approach) is relatively well supported or understood. Human disease and aberrant mutant phenotypes in laboratory model organisms are not included in Gephebase and can be found in other databases (*eg*. OMIM, OMIA, Monarch Initiative). Gephebase contains more than 1700 entries. Each entry corresponds to an allelic difference at a given gene and its associated phenotypic change(s) between two species or between two individuals of the same species, and is enriched with molecular details, taxonomic information, and bibliographic information. Users can easily browse entries for their topic of interest and perform searches at various levels, whether phenotypic, genetic, taxonomic or bibliographic (*eg*. transposable elements, cis-regulatory mutations, snakes, carotenoid content, an author name). Data can be searched using keywords and boolean operators and is exportable in spreadsheet format. This database allows to perform meta-analysis to extract general information and global trends about evolution, genetics, and the field of evolutionary genetics itself. Gephebase should also help breeders, conservationists and others to identify the most promising target genes for traits of interest, with potential applications such as crop improvement, parasite and pest control, bioconservation, and genetic diagnostic. It is freely available at www.gephebase.org.

## INTRODUCTION

Mutations form the raw bulk of heritable variation upon which traits evolve. Identifying the DNA sequence modifications that drive phenotypic changes is a primary goal of modern genetics, and could greatly improve our understanding of the mechanisms behind biodiversity and adaptation. However, this research program would be most successful if it reaches comparative capacity, for instance by allowing us to detect trends across the Tree of Life (1–3). Advances in genome sequencing and editing are accelerating the rate of discovery of the loci of evolution at a quick pace, making data integration increasingly challenging, and it is now crucial to develop a universal, single resource integrating this body of knowledge. As of today, compilations of genotype-phenotype relationships are available for a limited number of species in taxon-specific databases, for example OMIA for domesticated animals (4), OMIM for humans (5), TAIR for *Arabidopsis* (6), FlyBase for Drosophila (7), or the Monarch Initiative across the main laboratory animal model species (8, 9). To date, there are no databases that consolidate genotype-phenotype relationships related to natural evolutionary cases across all Eukaryotes. For example, evolutionary changes in tigers, butterflies, monkeyflowers, or any non-traditional model organism are lacking from existing genotype-phenotype databases, preventing comparative insights on the diversity and similarities of sequence modifications that fuel the generation of observable differences in the living world.

To fill this gap, we developed Gephebase, a manually curated database that gathers published data about the genes and the mutations responsible for evolutionary changes in all Eukaryotes (mostly animals, yeasts and plants) into a single website. The content of Gephebase was developed over the past 10 years, with previous versions of the dataset published as supplementary spreadsheet files associated to two review articles, which successively compiled 331 entries (1), and 1008 entries (2). These datasets have been used by various authors to highlight several trends regarding the genetic basis of natural variation. For example, based on these compilations it was found that the mutations responsible for long-term evolution have distinct properties than the mutations responsible for short-term evolution (1, 10), that certain types of mutations are more likely to be fixed than others during the course of evolution (11), that independent evolution of similar traits in distant lineages often involves mutations in the same orthologous gene (2), that current data are biased towards a limited number of model organisms (12), and that the cis-regulatory tinkering of signaling ligand genes is a recurring mode of morphological evolution (13).

We have now created an online version of the Gephebase database, accessible at www.gephebase.org, and we describe here its various features.

## SNAPSHOT SUMMARY

In short, Gephebase is a searchable, manually curated knowledge-base of the genetic bases of phenotypic variation. Each entry is a pair of alleles associated to a trait variation, be it naturally existing (inter-or intraspecific), selected by breeders (domestication), or occuring during a bout of experimental evolution in the lab. For instance, forward genetic studies have determined that independently derived null mutations of the *Oca2* gene have caused an amelanic phenotype in at least two subterranean populations of cavefish (14), and in a breed of corn snake that has been selected for the pet trade (15). A Gephebase search for the *Oca2* gene name reveals these findings, accessible in summary tables (**Fig. 1**) or in a more detailed output (entry view, and CSV spreadsheet format). Gephebase also indicates that some *Oca2* allelic variants have been identified by Genome-Wide Association Studies of pigment variation. Importantly, the focus of Gephebase is always on genetic variations that emerge naturally - it never includes laboratory variants that were generated by random or directed mutagenesis. Thus the *Oca2* CRISPR knockout phenotypes that have been generated in frogs (16) do not have a dedicated Gephebase entry; the cavefish *Oca2* CRISPR/TALEN knockout phenotypes (17, 18) do not have a dedicated entry either, but are used as Additional References to support the functionality of the two natural *Oca2* null alleles in Gephebase. This makes Gephebase complementary to the Monarch Initiative database, which compiles gene-to-phenotype relationships in humans, as well as in laboratory organisms and mutants generated by reverse genetics, but does not include non-model species such as cavefishes and corn snakes (8, 9).

**Fig. 1.**
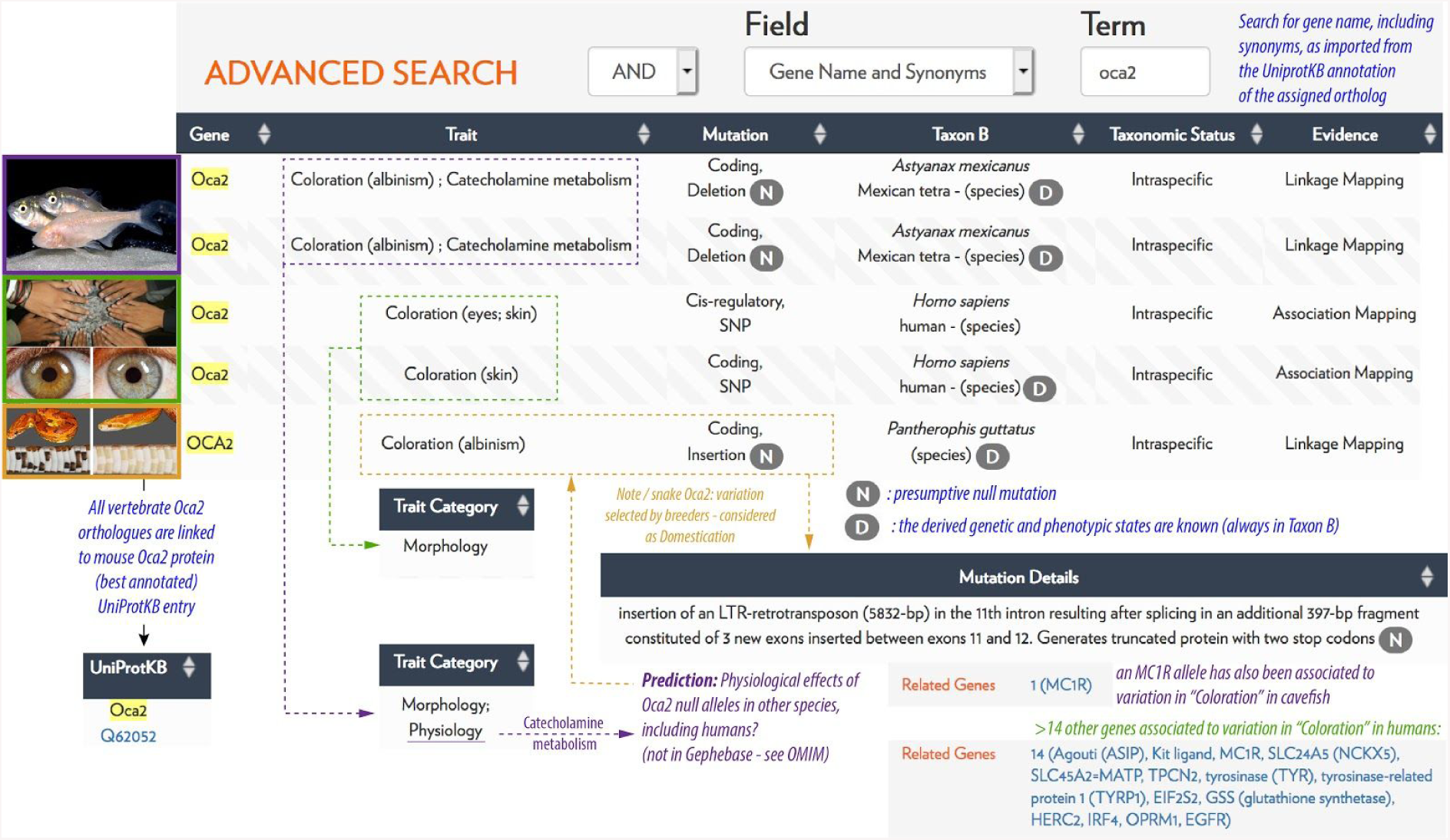
A snapshot view of the potential of Gephebase for comparative genetics. Part of the Summary Output Table of a gene name search for “oca2” is provided here as an example, with a montage of Gephebase summary outputs, pictures, and annotations in color. Interestingly, the cavefish studies reveal that the *Oca2* null alleles also affect catecholamine metabolism in cavefish. Gephebase can be used as a hypothesis generator: by juxtaposition of these entries, one may expect similar effects in the corn snake *Oca2* mutants that remain to be tested. “N” means that the mutation is null, “D” means that Taxon B is inferred to bear the derived trait. Photo Credits (top to bottom, with permission or license to reuse): Richard Borowsky, Flickr user *neokainpak* (CC BY 2.5), Aaron Pomerantz, Michel Milinkovitch.

## DATABASE CURATION AND STRUCTURE

### Criteria for inclusion in Gephebase

Gephebase includes cases of domestication, experimental evolution and natural evolution but no human clinical phenotypes. Gene expression levels (eQTL) and DNA methylation patterns are not included. All kinds of traits above this level, whether morphological, physiological or behavioral, are included. For example, we include “Recombination rate”, “Telomere length”, “Hematopoiesis”, “Hybrid incompatibility”.

Cases of genomic regions associated with a trait for which the underlying gene(s) is unclear are not included in Gephebase. Cases where the gene has been identified, but not the exact mutation, are included. Gephebase intends to compile studies where a given genotype-phenotype association is relatively well supported or understood. There are multiple types of experimental evidence that led to the discovery of a relationship between a genetic mutation and a phenotypic change. For sake of simplicity and efficiency, each gene-phenotype association is attributed only one type of Experimental Evidence among three possibilities: “Association Mapping”, “Linkage Mapping”, or “Candidate Gene” (**Fig. 2)**. When several methods were used, the least biased one is chosen by the curator (Table 1).. Gene-to-phenotype identified by Linkage Mapping with resolutions below 500kb have priority in the dataset. Association Mapping studies are included based on individual judgment, with a strong bias towards associations that have been confirmed in reverse genetics studies.

**Fig. 2.**
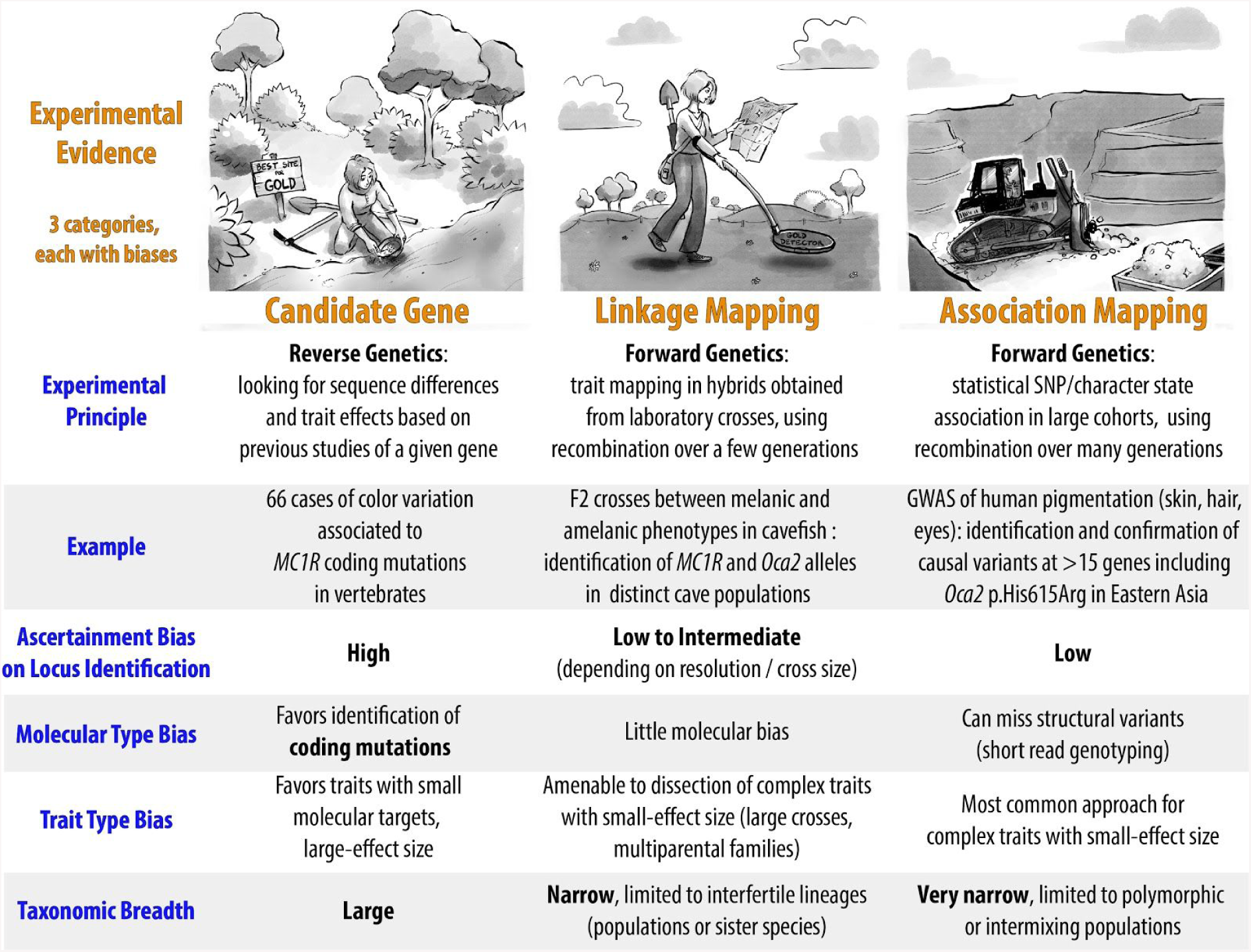
Three kinds of experimental strategies for identifying gene-to-phenotype variations. Gephebase focuses on genes that have been mapped using a forward genetics approach, and supported as the causal agents by sufficient evidence. Candidate gene approaches are also included and cover broader phylogenetic distances (*eg*. human/chimp), but tend to be biased towards the identification of coding changes for relatively simple traits. The search for the genes and mutations that drive phenotypic variation is somewhat analogous to searching gold: from left to right, targeted candidate gene approaches can identify variants of large effects at loci previously identified in other organisms; in a linkage mapping approach, the experimenter walks on chromosomes to narrow down the causal genetic interval, and can increase resolution and sensitivity with the analysis of more recombinants; association mapping (*eg*. GWAS) takes advantage of statistical power across large datasets to extract genetic variants in linkage disequilibrium with the causal mutations. GWAS: Genome-Wide Association Studies, SNP: Single Nucleotide Polymorphism.

**Table 1.**
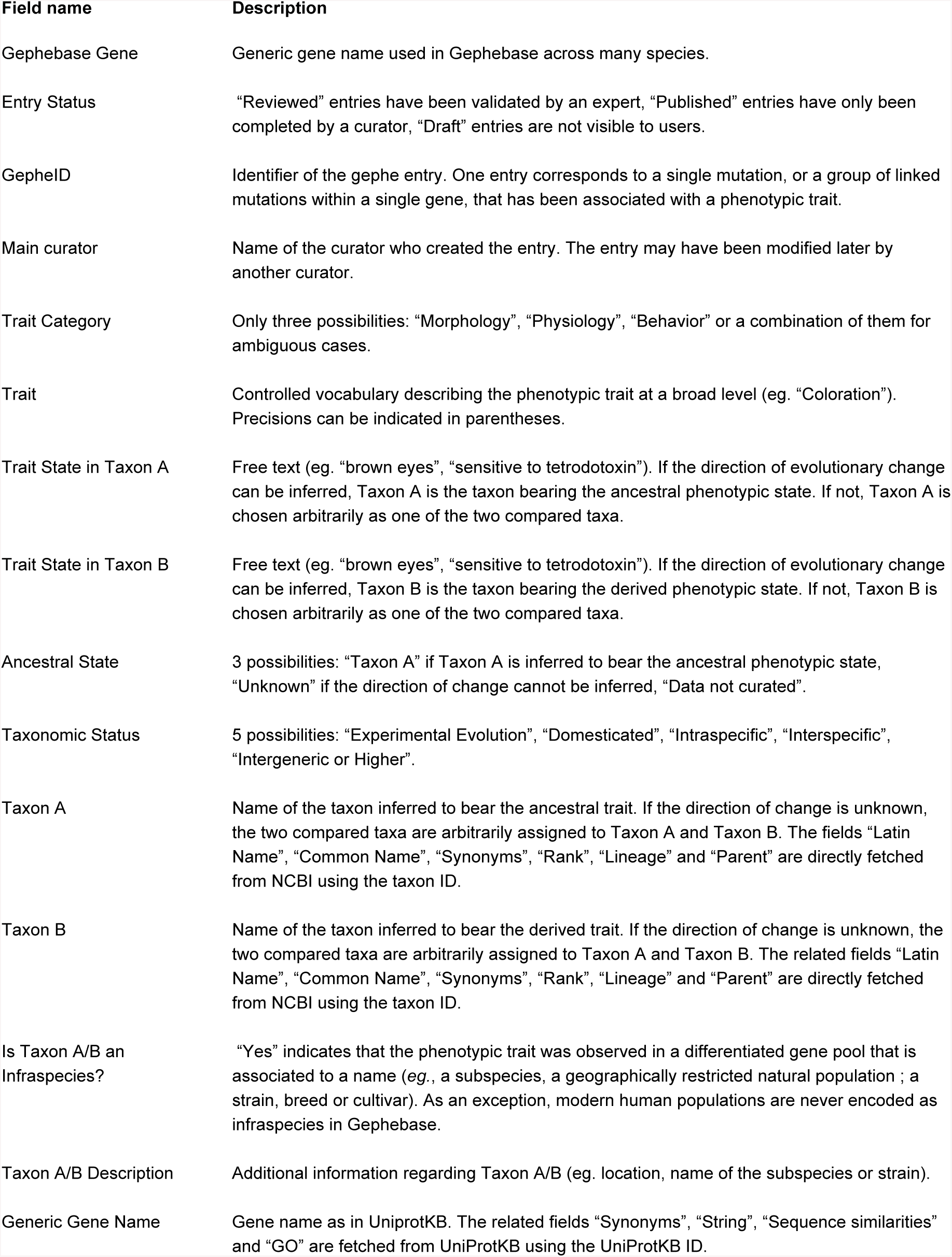
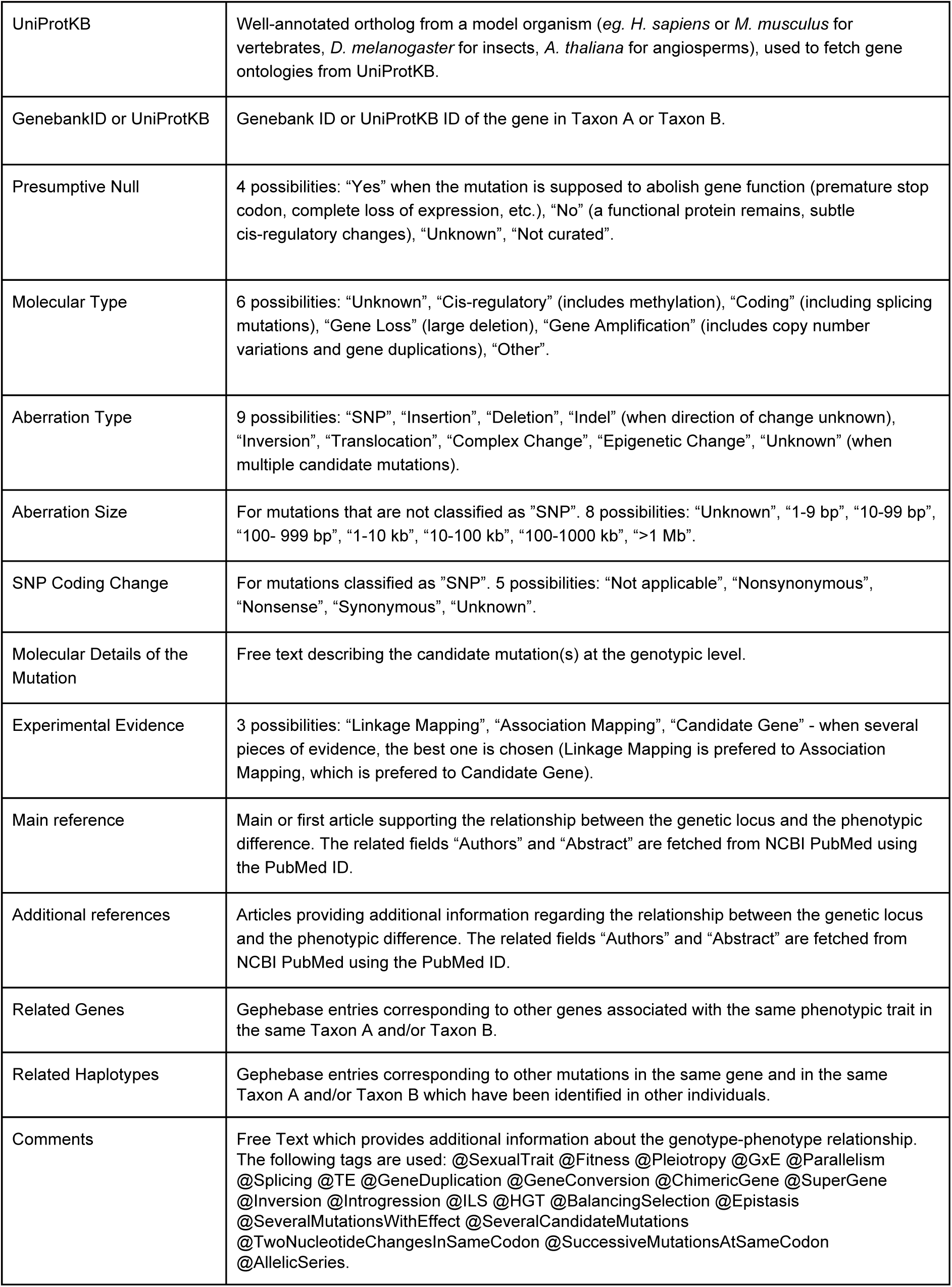
List of the fields of a Gephebase entry. (in order of appearance on the View-Entry page).

### Curation protocol

Searches for relevant papers to be included in the database are done manually by our team of curators. We screen major journals in evolutionary genetics, perform keyword searches using online search tools, and we pay particular attention to citations in primary research articles as well as in review papers. The “Suggest an article” button in the top bar menu allows users to suggest articles to our curation team. We believe that the database is rather comprehensive until 2013 for all species.

### Technical overview of the database and the web interface

Gephebase was developed using the Symfony framework (v2.8) and PHP (v5.6 compatible 7). MariaDB (v10) is used to store data. The database consists of 33 tables including users management and logs. The main table links genotypic change, phenotypic change, references and validation information. Most fields of other tables are automatically retrieved from NCBI databases. The import procedure uses the NCBI E-utility interface with XML to fill the corresponding tables. Gephebase entries of the main table can be imported and exported through a csv file. For convenience, fields retrieved automatically can be present in the csv file even though they are fetched and are stored in other tables.

The project code was put under version control (git) from its inception. The code will be released in the GitHub repository https://github.com/Biol4Ever/Gephebase-database under GPL (GNU General Public License) version 3. The content of the database (Table S1) will also be deposited in GitHub at https://github.com/Biol4Ever/Gephebase-database.

## DATABASE CONTENT AND WEB INTERFACE

### Organization of the data into entries

This database currently comprises more than 1700 entries. One entry corresponds to a single mutation, or a group of linked mutations within a single gene, either between two closely related species or between two individuals of the same species, and its associated phenotypic change **(Fig. 3)**. For cases of repeated evolution (2), we use the following conventions. When several mutations are found within the same gene in a given individual, with each mutation affecting the trait of interest - *ie*. several causative mutations within a haplotype, intralineage hotspot (2)-all are grouped into a single entry. In contrast, when independent mutations occur in the same gene in distinct individuals of the same species, leading to similar phenotypic changes (intraspecific parallel evolution, convergent evolution), we chose to create different entries for each lineage-specific haplotype. In cases where a genetic variant was invented once, and then spread into multiple branches of the gene pool, via Incomplete Lineage Sorting (ILS), secondary hybridization (introgression among organisms that are not completely reproductively isolated) or horizontal transfer, a single entry is created and multiple taxa with the derived trait are reported in the entry.

**Fig. 3.**
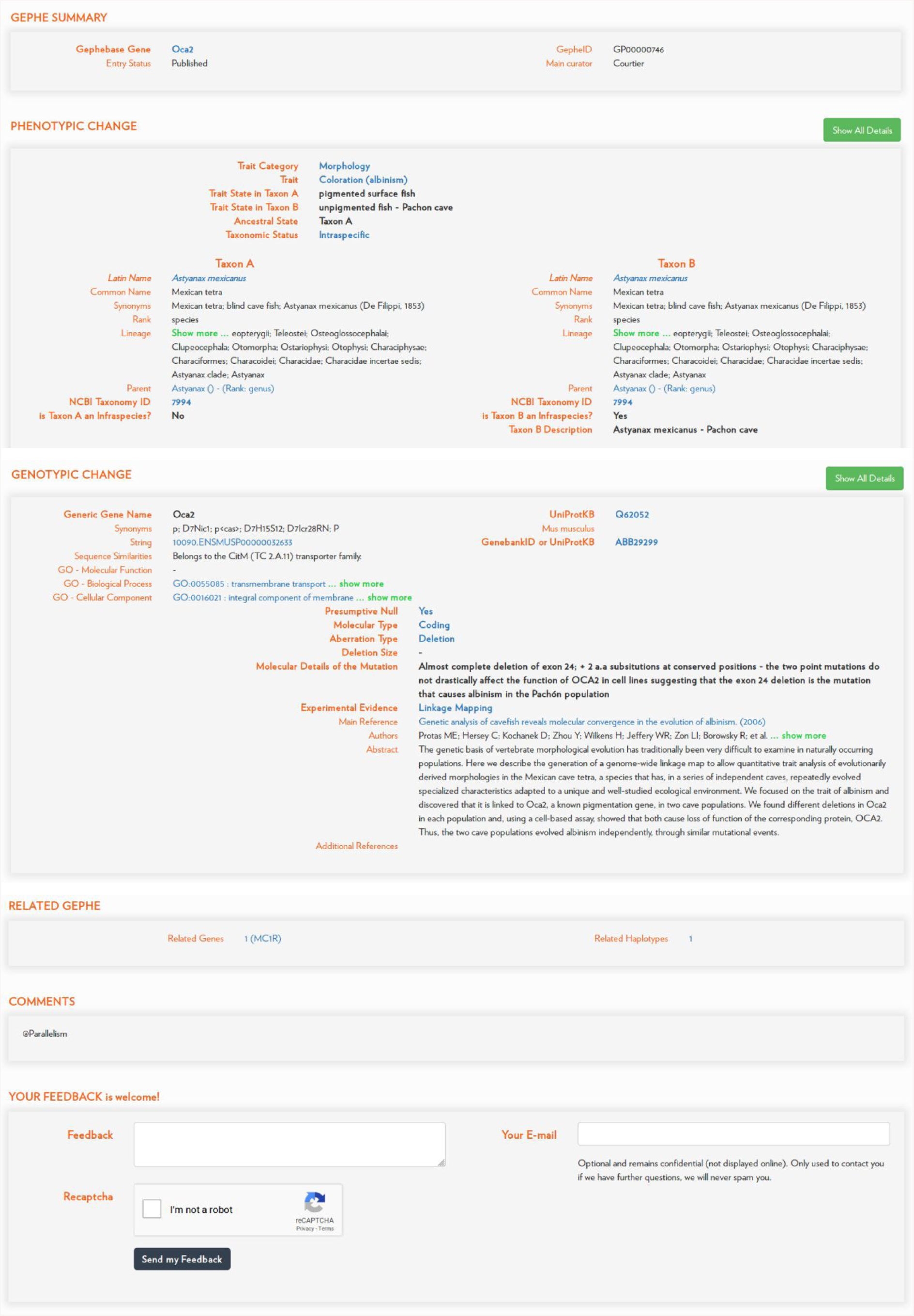
Example of a Gephebase entry. Gephe entries provide extensive information about phenotypic traits, taxon groups, genetic changes, as well as publications.

### The various fields of a Gephebase entry

A Gephebase entry (**Fig. 3**) comprises 29 manually curated fields regarding bibliographical information, molecular details and taxonomic information; some are free-text and others rely on controlled vocabulary (Table 1). In addition, for each entry, 20 fields are automatically fetched based on manually curated data, from NCBI Taxonomy using the Taxon ID (17), from UniProtKB using the UniProtKB ID (18), and from NCBI PubMed using the PubMed ID (19). Two fields are also automatically computed within Gephebase: “Related Genes”, which corresponds to the other genes in Gephebase associated with the same phenotypic trait in the same group of species, and “Related Haplotypes”, which displays the other mutations in Gephebase that are found in the same gene and that occurred in other lineage branches in the same group of species.

A single entry can include several traits if a mutation is pleiotropic. Taxon A represents the taxon(s) inferred to bear the ancestral phenotypic state and Taxon B the derived state. If the direction of change cannot be inferred, the field “Ancestral State” is “Unknown” and the two compared taxa are assigned arbitrarily to Taxon A and Taxon B. In most cases, Taxon A and Taxon B correspond to taxa at the species level. In cases of named breeds, cultivars, strains or geographically restricted populations, additional information about the Taxon A/B can be found in the field “Taxon A/B Description”. The phenotypic states are described in “Trait State in Taxon A/B”.

### Exploration tools

Gephebase is designed for interactive exploration and analysis of the genotype-phenotype relationships across species and populations. First-time users can find help on the Frequently Asked Questions page, in tutorials available on the Documentation page and via “contextual tips”, small boxes providing information when the cursor hovers over an item. Data can be queried using boolean operators via the Search page, via SQL line or via custom tools after downloading the dataset of interest as a csv file. The entire dataset can be downloaded as a CSV file by searching for the wild card * in the top bar panel, clicking on “Select all” in the top left corner of the results table and then clicking on “Complete Export”.

Gephebase comprises two main views, the View-Entry page which displays a single entry (**Fig. 3**) and the Search/Results page, which shows the results of a given search in a table format (**Fig. 4**). Results of a search are displayed as a table and there are four view options. In the default view, one line of the table corresponds to one Gephebase entry. Under the option “Split Mutations”, one line corresponds to one mutation. Under the option “Group Haplotypes”, one line corresponds to all haplotypes of a given gene for a given pair of Taxon A and Taxon B. Under the option “Group Genes”, one line corresponds to all the genes associated with the same phenotypic trait in the same Taxon A and Taxon B. The number of lines in the Results Table is indicated above the table (for example “10 results” in **Fig. 4**). Clicking on one line of the Results Table will display all the corresponding Gephebase entries if the line corresponds to several Gephebase entries and will lead to the corresponding View-Entry page if the line corresponds to one entry.

**Fig. 4.**
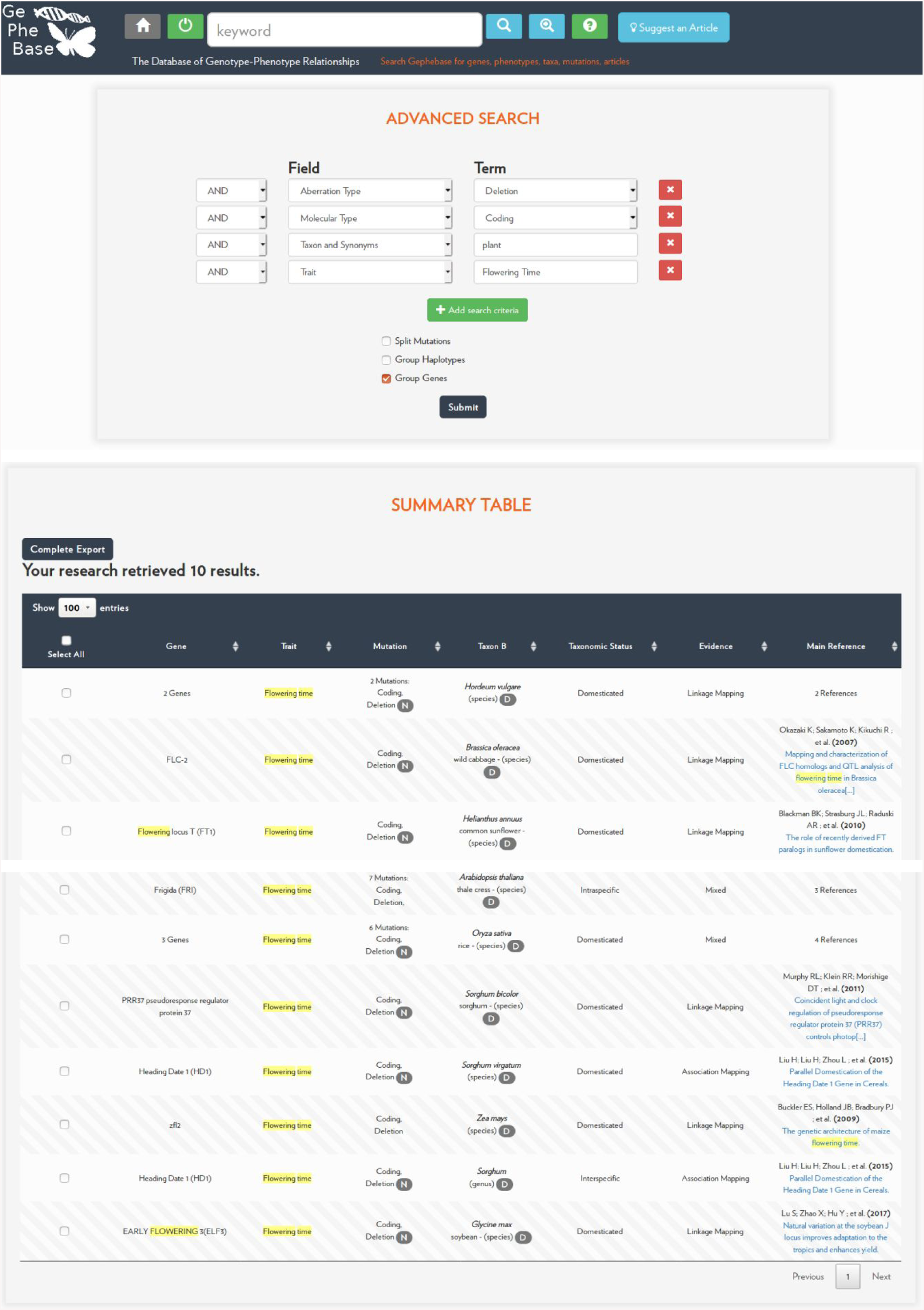
Results of an advanced search. In this example, the upper panel shows a query form for “Coding” “Deletion” in “plant” affecting “Flowering time” and below is the Results Table. The option “Group Genes” bundles the entries together when they match the same species and trait. “N” means that the mutation is null, “D” means that Taxon B is inferred to bear the derived trait.

Extensive links to external databases (UniProtKB, NCBI Taxonomy, NCBI PubMed) and to Gephebase itself allow in-depth analysis of curated data. Users can provide feedback using the Feedback section on each View-Entry Page (**Fig. 3**) and can suggest new articles for curation in Gephebase using the “Suggest an article” button in the top bar menu.

## DISCUSSION AND CONCLUSION

Gephebase contains data for more than 420 eukaryote species and more than 890 distinct genes. In Gephebase, physiological traits represent 64% of the entries, morphological traits 34%, behavioral traits 1% and mixed morphological/physiological/behavioral traits 1% (**Fig. 5A,C**). The most represented traits are Coloration (19% of the entries) and Xenobiotic Resistance (19%). The most represented taxa are Vertebrates (40% of the entries), Green Plants (31%) and Arthropods (16%) (**Fig. 5D**), and a large contribution from traditional model organisms (12). Most data correspond to intraspecific changes (48% of the entries) and domesticated cases (30%) whereas interspecific cases correspond to 11.5% of the entries (**Fig. 5E**). The three categories of Experimental Evidence are relatively well-distributed among entries (**Fig. 5F**). Gephebase contains a higher number of coding mutations (63% of the entries) compared to cis-regulatory changes (18% of the entries, **Fig. 5G**). While a significant fraction of Gephebase correspond to cases where the exact mutation has not been identified (24% of entries with Aberration Type “Unknown”, **Fig. 5H**), most mapped mutations are single nucleotide changes (47%) and indels (23%).

**Fig. 5.**
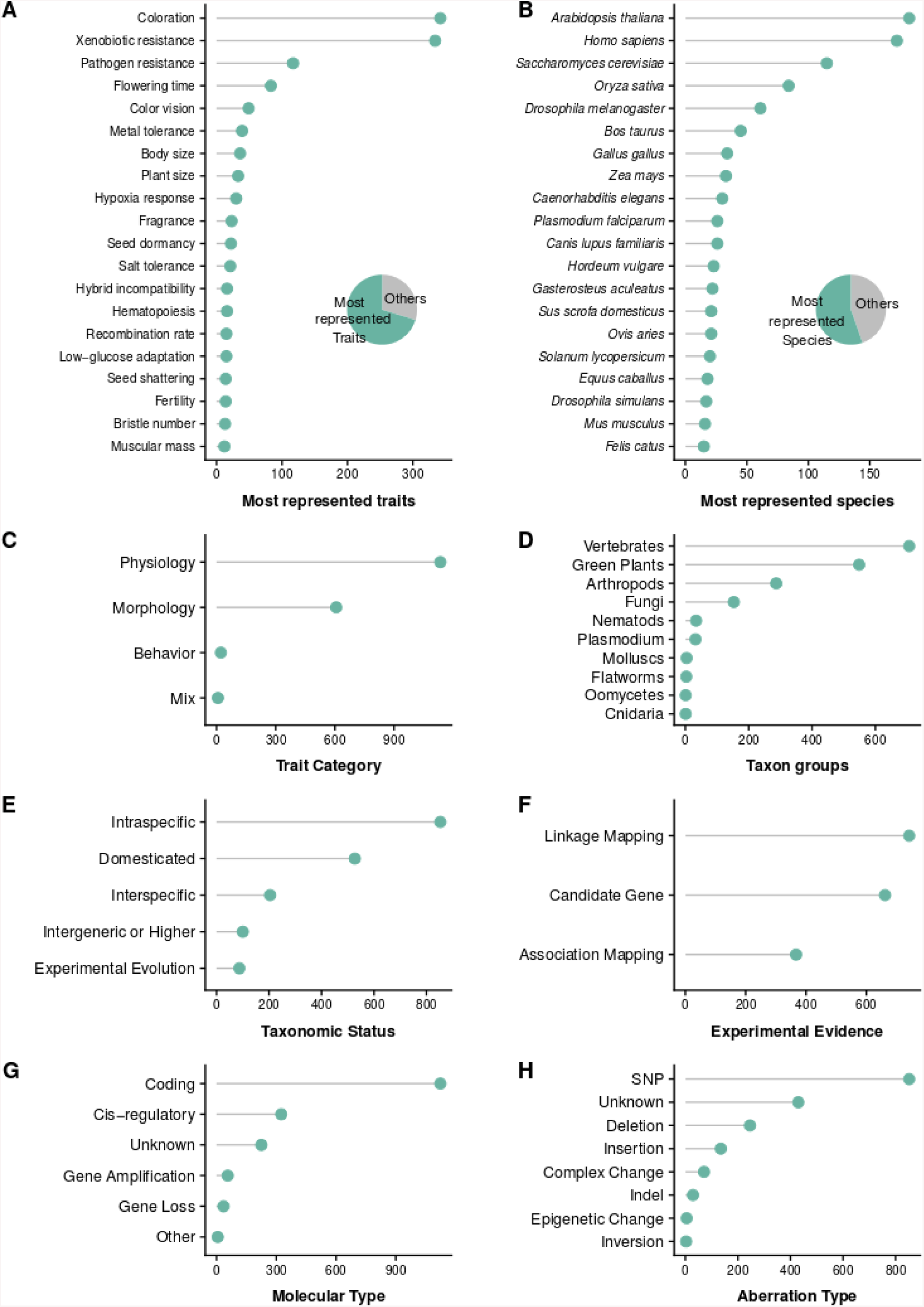
Summary statistics for current Gephebase data. Y-axes indicate the number of entries for each category. (A) Distribution of the twenty most represented Traits. (B) Distribution of the twenty most represented Taxon B species. Pie charts in (A-B) show the distribution of all entries between the twenty most represented Traits (A)/Species (B) and the remaining others. (C) Distribution of Trait Categories (Morphology, Physiology, Behavior, Combination of at least two trait categories). (D) Distribution of Taxon Groups. (E) Distribution of Taxonomic Status. (F) Distribution of Empirical Evidences. (G) Distribution of Molecular Types of genetic changes. “Other” corresponds to mutations creating chimeric genes or to super gene loci. (H) Distribution of Aberration Types. “Indel” corresponds to cases where the direction of change is unknown. When several mutations are reported within one entry, only the first curated mutation of the entry was used for statistical analysis.

Gephebase stands out compared to the other current databases of genotype-phenotype relationships in that it compiles genotype-phenotype data across all Eukaryotes. We consider our dataset to be highly complementary to other available databases, which are more species-specific and which usually include more detailed information about genotype-phenotype relationships. Gephebase is of interest not only to researchers working on the genetic basis of phenotypic variation, but also to breeders interested in transferring traits of interest to new species, and to epistemologists interested in biases and sociological aspects in the field of genetic evolution.

## Supporting information

Full content of Gephebase

R script used to create Fig. 5

## DATA AVAILABILITY

Gephebase is freely available at gephebase.org. The code will be available on GitHub. The entire dataset is freely available for download by searching for “*” and clicking on “Complete Export”.

## SUPPLEMENTARY DATA

Table S1. Complete Gephebase dataset as of April 2019 (1771 entries).

Data S1. R script used to create Fig. 5.

## ACKNOWLEDGEMENTS

We thank the twenty participants of the “Loci Of Evolution Workshop” (Paris, September 2016) for their enthusiasm and encouragements. We are indebted to the team of AtoutLibre (France) and especially Kyle Ratteree for developing the software and website behind Gephebase. Nathalie Vessilier drew the illustrations featured in Figure 2.

